# Genetic mapping of fitness determinants across the malaria parasite *Plasmodium falciparum* life cycle

**DOI:** 10.1101/570085

**Authors:** Xue Li, Sudhir Kumar, Marina McDew-White, Meseret Haile, Ian H. Cheeseman, Scott Emrich, Katie Button-Simons, François Nosten, Stefan H.I. Kappe, Michael T. Ferdig, Tim J.C. Anderson, Ashley M. Vaughan

## Abstract

Malaria is transmitted through female Anopheline mosquitoes where gamete fusion and meiosis occurs, and humans where parasites proliferate asexually. We describe a powerful approach to identify the genetic determinants of parasite fitness across both invertebrate and vertebrate life-cycle stages in human malaria parasite *Plasmodium falciparum* using bulk segregant analysis (BSA). We combined experimental genetic crosses using humanized mice, with selective whole genome amplification and BSA at multiple developmental stages in both mosquito and vertebrate host to examine parasite competition and identify genomic regions under selection. We generated crosses between artemisinin resistant (ART-R, *kelch13*-C580Y) and ART-sensitive (ART-S, *kelch13*-WT) parasite clones recently isolated from Southeast Asian patients. We then quantified genome-wide changes in allele frequency in the parasite progeny population from infected midgut and salivary glands of *Anopheles stephensi* mosquitoes, infected livers, emerging merozoites and aliquots of *in vitro* cultured progeny parasites at intervals over 30 days. Three striking results emerge: we observed (i) a strong skew (>80%) towards alleles from the ART-R parent in the mosquito stage, that dropped to ∼50% in the blood stage as selfed ART-R parasites were selected against; (ii) highly repeatable skews in allele frequencies across the genome in blood stage parasites; (iii) particularly strong selection (selection coefficient (s) ≤ 0.18/asexual cycle) against alleles from the ART-R parent at loci on chromosome 12 containing MRP2 and chromosome 14 containing ARPS10. This approach robustly identifies selected loci and has strong potential for identifying parasite genes that interact with the mosquito vector or compensatory loci involved in drug resistance.

## Introduction

Parasitic organisms frequently use multiple hosts and have several morphologically and transcriptionally distinctive life cycle stages. Within each host, parasites must circumvent immune defenses and navigate to new tissues. There are frequently extreme bottlenecks in parasite numbers during transmission (Hopp et al. 2015), with rapid proliferative growth within hosts, and intense competition between co-infecting parasite genotypes. For example, the life cycle of malaria parasites involves successive infection of two hosts: female *Anopheles* mosquitoes, where gamete fusion, meiosis and recombination occurs, and humans in which parasites travel from the skin, develop in the liver and then proliferate asexually in the blood stream. Ideally, we would like to understand how natural selection operates across the complete life cycle and document the genes subject to selection pressures at each life cycle stage: during erythrocytic growth, gametocyte production, oocyst development in the mosquito midgut, migration of sporozoites to the salivary glands, transmission from the salivary glands, sporozoite survival in the skin, and establishment and parasite growth during liver stage development and exoerythrocytic merozoite release.

Selection can be directly measured by examining changes in allele frequency across these developmental stages. Shifts in allele frequencies in populations of thousands of progeny generated by experimental genetic crosses provide locus-specific readouts of competitive fitness. For example, deep sequencing of bulk populations containing thousands of recombinants identified yeast genes selected under different regimens (Ehrenreich et al. 2010; Parts et al. 2011; Feng et al. 2018). Bulk segregant analysis (BSA) has also been successfully applied to studies of several different parasitic organisms including coccidia (*Eimeria tenella*) and the human blood fluke *Schistosoma mansoni* (Blake et al. 2011; Chevalier et al. 2014). Our work was inspired by an exciting series of papers applying pooled sequencing approaches (termed linkage group selection in the malaria literature) for mapping genes of interest in rodent malaria parasites (Rosario et al. 1978; Culleton et al. 2005; Martinelli et al. 2005; Pattaradilokrat et al. 2009; Hunt et al. 2010).

Most studies of *Plasmodium falciparum* to date focus only on the asexual erythrocytic stages (Rosario et al. 1978; Walliker et al. 2005; Petersen et al. 2015; Straimer et al. 2017; Nair et al. 2018), because they can be easily cultured *in vitro* in red blood cells, circumventing the need for humans or great apes, the natural hosts for this parasite. Two new research tools now allow us to examine selection across the complete life cycle of *P. falciparum*. First, we can maintain the complete life cycle of *P. falciparum* in a laboratory setting by using humanized mice (Vaughan et al. 2015) in place of splenectomized chimpanzees or human volunteers. These mice contain human hepatocytes and are therefore able to support liver stage development of *P. falciparum*. Hence, we can stage genetic crosses between different *P. falciparum* parasites, including parasites recently isolated from infected patients, and sample multiple parasite life stages for measurement of allele frequency changes throughout the life cycle. Second, selective whole genome amplification (sWGA) provides a simple and effective way to enrich *Plasmodium* DNA from contaminating host tissues. This is critical because *Plasmodium* DNA constitutes a very small fraction of DNA present in malaria-infected mosquitoes; likewise, *Plasmodium* DNA makes up a very small fraction of DNA extracted from malaria-infected livers (Table 1). sWGA uses short 8-12 mer oligonucleotide probes that preferentially bind to the target genome, rather than random hexamers used in normal whole genome amplification. This approach was pioneered by Leichty and Brisson (Leichty and Brisson 2014), and protocols for sWGA have been successfully developed to amplify and sequence malaria parasite genomes from contaminating host tissues (Guggisberg et al. 2016; Oyola et al. 2016; Sundararaman et al. 2016; Cowell et al. 2017).

**Table 1.** Sample collection and sequence statistics.

| Parasite stage | (1) Early Oocyst | (2) Maturing oocyst | (3) Sporozoite | (4) Liver Stage | (5) *In vivo* Blood | (6) *In vitro* Blood |
| --- | --- | --- | --- | --- | --- | --- |
| Collecting time<sup>a</sup> | d4 | d10 | d14 | d21 | d21 | d22-52 |
| Sample collected | 48 midguts | 48 midguts | 200 Salivary glands | 60 mg liver | 50ul blood (3.5% parasitaemia) | 50ul blood (1-4% parasitaemia) |
| Total DNA (ng) | 1,397 | 1,337 | 3,675 | 9,359 | 142 | 154-2,576 |
| Total *P. falciparum* genome copies<sup>b</sup> | 7,563 | 866,299 | 4,726,149 | 12,521,577 | 1,246,535 | 19.1M-279.6M |
| *P. falciparum* DNA percent before sWGA | 0.01% | 1.80% | 3.21% | 3.24% | 30.32% | 100% |
| Sequencing approach<sup>c</sup> | Amplicon | Amplicon, sWGA-WGS | Amplicon, sWGA-WGS | Amplicon, sWGA-WGS | Amplicon, sWGA-WGS, WGS | Amplicon, sWGA-WGS, WGS |
| Copy of *P. falciparum* genome for sWGA | na | 2×10<sup>5</sup> | 2×10<sup>5</sup> | 2×10<sup>5</sup> | 2×10<sup>5</sup> | 2×10<sup>5</sup> |
| *P. falciparum* DNA percent after sWGA<sup>d</sup> | na | 88.09% | 86.74% | 97.16% | 95.31% | 97.33%-99.57% |

b, We qualified the parasite genome copy number in the total DNA using qPCR, and translated this into parasite DNA percentage, using 2.48×10<sup>-5</sup> ng as the weight of *Plasmodium* genome.
c, For samples with host contamination or small amounts of DNA isolated (parasite stages 1-6), we performed selective whole genome
amplification (sWGA) before whole-genome sequencing (WGS). We used amplicon sequencing to trace biases in mtDNA transmission in those samples. For *in vitro* blood samples, we performed sequencing both before and after sWGA to evaluate the accuracy of allele frequency estimated after sWGA. To get sufficient representation of the bulk segregant samples, we used $2 \times 10^5$ copies of parasite genome as template for each sWGA reaction and 1,000 copies for amplicon sequencing. d, *P. falciparum* DNA percentage after sWGA was measured as the percent of reads that mapped to the *P. falciparum* 3D7 genome.

Artemisinin resistance is currently spreading across Southeast Asia (Ariey et al. 2014). SNPs in *Kelch13* (PF3D7_1343700) locus on chromosome (chr) 13 underlie resistance and greater than 124 independent alleles have been recorded in dramatic example of a soft select sweep (Anderson et al. 2016; Fairhurst and Dondorp 2016). One particular allele (*Kelch13*-C580Y) is currently replacing other resistant alleles and spreading toward fixation in independent transmission foci in western Cambodia/Laos/Vietnam and the Thailand-Myanmar border (Takala-Harrison et al. 2014; MalariaGEN Plasmodium falciparum Community Project 2016; Imwong et al. 2017). Several studies have suggested that mutations within loci other than *Kelch13* may provide a permissive background for evolution of artemisinin resistance or play a compensatory role (Miotto et al. 2015; Cerqueira et al. 2017), but the role of such accessory loci is poorly understood.

In this study, we measured skews in allele frequencies across the genome in the progeny of a genetic cross between artemisinin resistant (ART-R, *kelch13-*C580Y) and ART sensitive (ART-S, *kelch13*-WT) parasites throughout the life cycle to identify genes that influence parasite fitness in parasite stages infecting both the mosquito and vertebrate host. We selected ART-R and ART-S parental parasites in order to examine loci contributing to fitness and compensation for deleterious effects of ART-R alleles (Nair et al. 2018). We used the humanized mouse model to allow parasite liver stage development of the genetic cross progeny, sWGA to enrich parasite DNA from host contaminations and pooled sequencing to determine temporal change in allele frequency and characterize genomic regions under selection. Our results demonstrate pervasive selection across the parasite genome over the course of a single parasite generation, selection against progeny produced from selfed matings, and strong locus-specific selection against parasite loci on chr 12 and 14.

## Results

### Identification of high-confidence SNPs between parents

*P. falciparum* NHP1337 and MKK2835 were cloned by limiting dilution and used as parents for genetic crosses. MKK2835 (ART-S) is a *kelch13* wild-type ART-susceptible parasite collected from a patient visited the clinic in 2003 before ACT therapy was used. NHP1337 is a recent cloned ART-R parasite, that cleared slowly (Clearance half-life (T_½_P) = 7.84 h) from the blood of a patient treated with artemisinin combination therapy and carries the C580Y *kelch13* mutant. Parasites with C580Y mutation have been rapidly spreading in Southeast Asia and are replacing other ART-resistant *kelch13* alleles (Ashley et al. 2014; Anderson et al. 2016). We sequenced both parental parasites with coverage > 100×. To reduce false positives due to alignment errors, we excluded the high variable genome regions (subtelomeric repeats, hypervariable regions and centromeres) and only performed genotype calling in the 21 Mb core genome (defined in (Miles et al. 2016)). After filtering, we detected 9,462 high confidence SNPs between the two parental strains. These SNPs are distributed across the genome, with an average of 1 SNP per 2.43kb (Supplemental Fig. S1).

### Genetic cross and generation of segregant pools

To generate segregant pools of progeny, we crossed NHP1337 and MKK2835 (Fig. 1). We fed 500 mosquitoes with a ∼50:50 gametocyte mixture of the two parental parasites. Recombinant progeny are generated after gametes fuse to form a zygotes that then rapidly transforms into a short-lived tetraploid ookinetes which migrates to the basal lamina of the mosquito midgut and transforms into an oocyst. Mitotic division of the 4 meiotic products ultimately leads to the generation of ∼10,000 haploid sporozoites within each oocyst. Oocyst prevalence was 80% with an average burden of three oocysts per mosquito midgut (range: 0-6), giving an estimate of 12 (3×4) recombinant genotypes per mosquito. We dissected a proportion of the infected mosquitoes to collect midguts (48 at each time point) for monitoring allele frequencies during oocyst development. Salivary gland sporozoites from 204 mosquitoes were pooled together and injected in to a single FRG huHep mouse, which resulted in an estimate of 2448 (204 × 12) recombinants in this inoculation.

**Figure 1.**
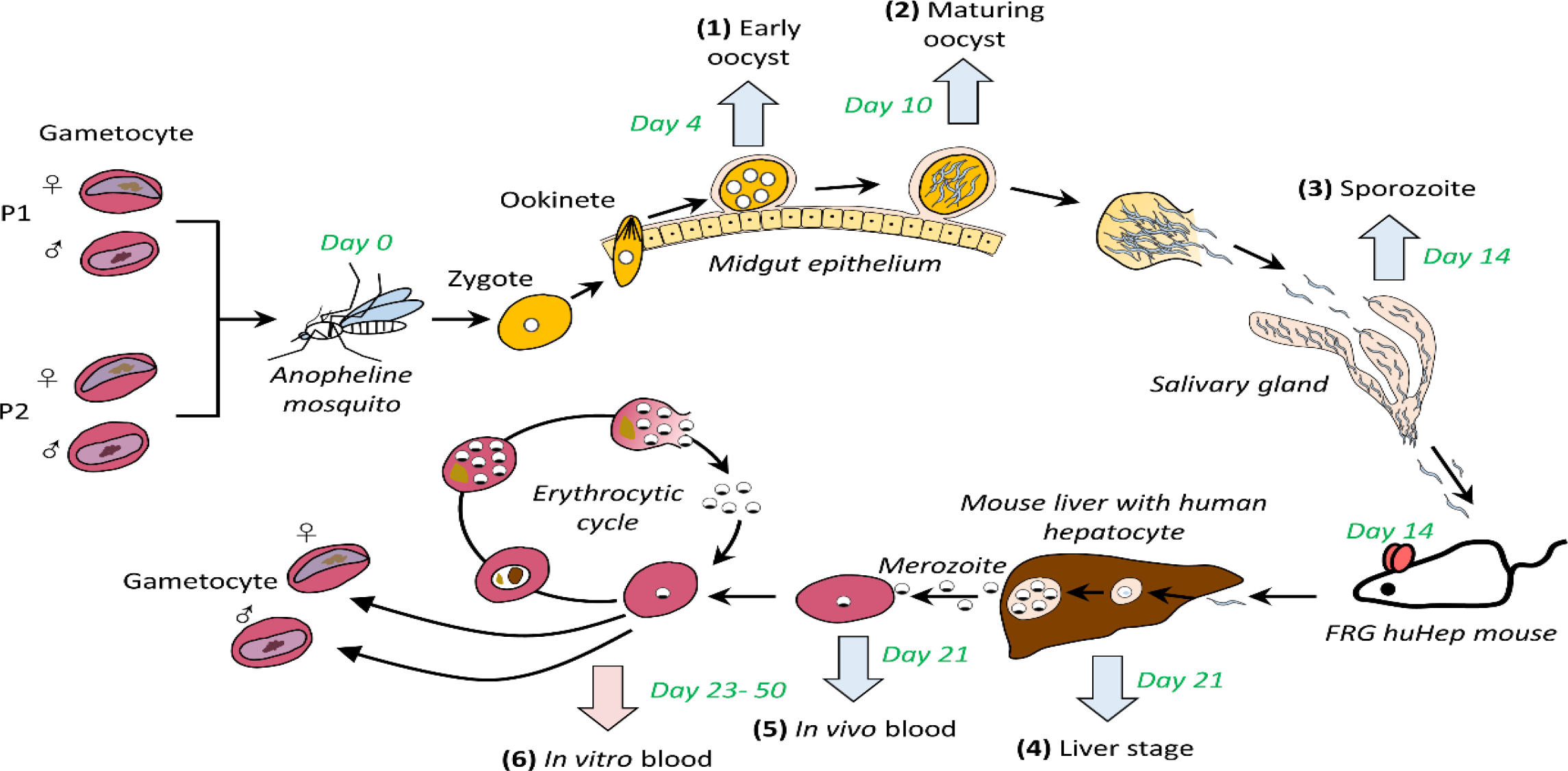
Genetic mapping of parasite competition through *Plasmodium falciparum* life cycle. We generated genetic crosses using *Anopheles stephensi* mosquitoes and FRG huHep mice. We collected midgut and salivary glands from infected mosquitoes, infected mouse liver and emerging merozoites from *in vivo* blood, and recovered aliquots of *in vitro* cultured progeny parasites at intervals of 30 days (marked with arrows, parasite stage 1-6). Cross generation and sample collection was completed in two months (marked with green letters). For samples with host contamination or small amounts of DNA isolated (blue arrows, Table 1), selective whole genome amplification (sWGA) was performed before Illumina whole-genome sequencing (WGA). We used amplicon sequencing to trace biases in mtDNA transmission in those samples. For *in vitro* blood samples (pink arrow), we performed sequencing both before and after sWGA to evaluate the accuracy of allele frequency after sWGA.

We collected samples for allele frequency analysis from infected mosquito midguts, infected mosquito salivary glands, infected humanized mouse livers and infected blood (both mouse blood and injected human red blood cells) after the liver stage-to-blood stage transition. We then recovered aliquots of *in vitro* cultured progeny parasites at two-four day intervals over 30 days (Fig. 1). These samples represent the important developmental stages across the parasite life cycle, including early oocyst, maturing oocyst, sporozoites, liver stage schizonts, transitioned blood stage parasites and fifteen asexual cycles in blood stage culture (Table 1).

We measured the total number of parasite genome copies and the amount of host DNA contamination for these segregant pools using qPCR. At the early midgut oocyst stage (4 days after mosquito infection), we isolated ∼8,000 copies of the *P. falciparum* genome from 48 mosquito midguts. The parasite DNA represented approximately 0.01% of the total DNA within these isolated midguts. The percentage reached 1.80% after 10 days of mosquito infection, which indicating a 196-fold increase of parasite DNA in the six days following initial midgut isolation. The percentage of parasite DNA found in samples from mosquito salivary gland containing sporozoites, liver containing liver stage parasites and liver stage-to-blood stage transitioned *in vivo* blood samples were 3.21%, 3.24 % and 30.32%, respectively (Table 1).

### sWGA-WGS, WGS and amplicon sequencing

We used three approaches to sequence the segregant pools and quantify allele frequencies: (1) selective whole genome amplification combined with whole genome sequencing (sWGA-WGS), (2) direct whole genome sequencing (WGS) and (3) amplicon sequencing. The methods used were dependent on the level of host contamination and the total amount of DNA present in the samples (Table 1). We used multiple methods where possible to determine potential bias. We used the sWGA approach to enrich parasite DNA before WGS in samples with extensive host contamination, including the mosquito midgut and the FRG NOD human-chimeric mouse liver (Table 1, sample 2-5). With 0.2×10^6^ copies of parasite genome as template, the sWGA-WGS approach yielded 0.6-1.4 µg of product after 3h of amplification, of which > 88% was from *P. falciparum*, for both mosquito and mouse samples. By sequencing pools to ∼100× coverage, comparable results were obtained between samples prepared by the sWGA-WGS approach and the WGS approach (Fig. 2A; Fig. 3; Fig. 4). We used amplicon sequencing (Nair et al. 2018) to determine the frequencies of mtDNA from the two parents in those samples for which we used sWGA (Fig. 2C; Supplemental Fig. S2). This was necessary because our sWGA primers were specifically designed to minimize amplification of mtDNA, since we were concerned that sWGA with circular DNA would inundate autosomal sWGA products. For day 4 mosquito midgut samples, we only have amplicon sequencing data since there was insufficient parasite DNA for a successful sWGA.

**Figure 2.**
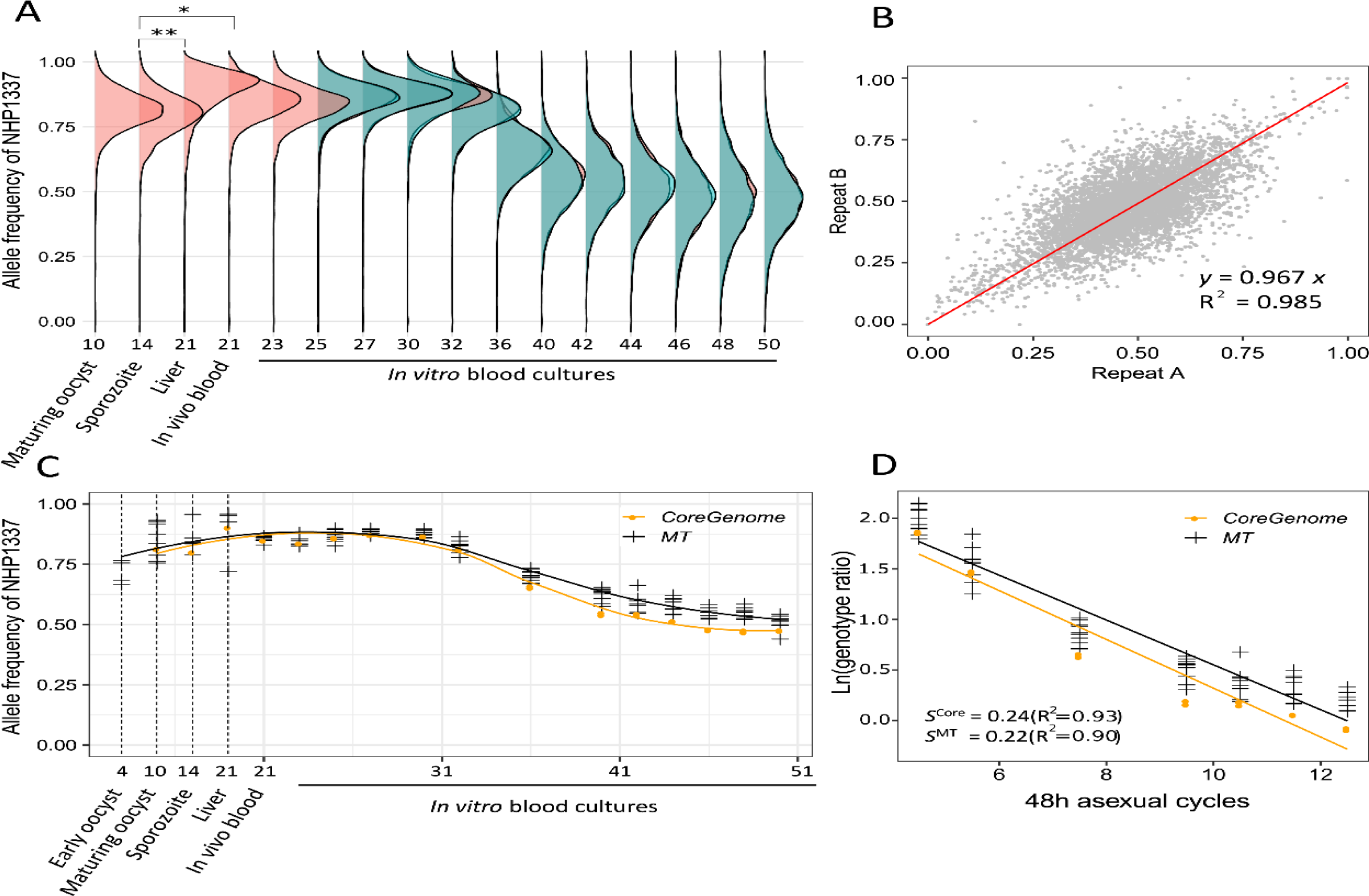
Change in frequency of mitochondria and core genome at different infection stages. (A) Ridgeline plots showed genome-wide allele frequency distributions of NHP1337 through the *Plasmodium* life cycle. * indicates Cohen’s *d* effect size > 0.5, and ** indicates effect size > 0.8. (B) We detected strong concordance between allele frequencies estimated from experimental replicates. (C) The allele frequency estimated from mitochondria and core genome showed the same pattern of skew with across the life cycle. (D) Natural log of the genotype ratio (NHP1337/MKK2835) plotted against asexual life cycles. The selection coefficient was estimated as the slope of the least-squares fit. Allele frequencies from day30 to day42 were used here. There was no significant difference between fitness costs estimated for the core genome and mitochondria (*P* = 0.363). Positive values of *s* indicate a selection disadvantage for NHP1337. MT, mitochondria; *s*, selection coefficients; R, correlation coefficient. X-axis in (A) and (C) indicated sample collecting days and corresponding parasite developmental stages.

**Figure 3.**
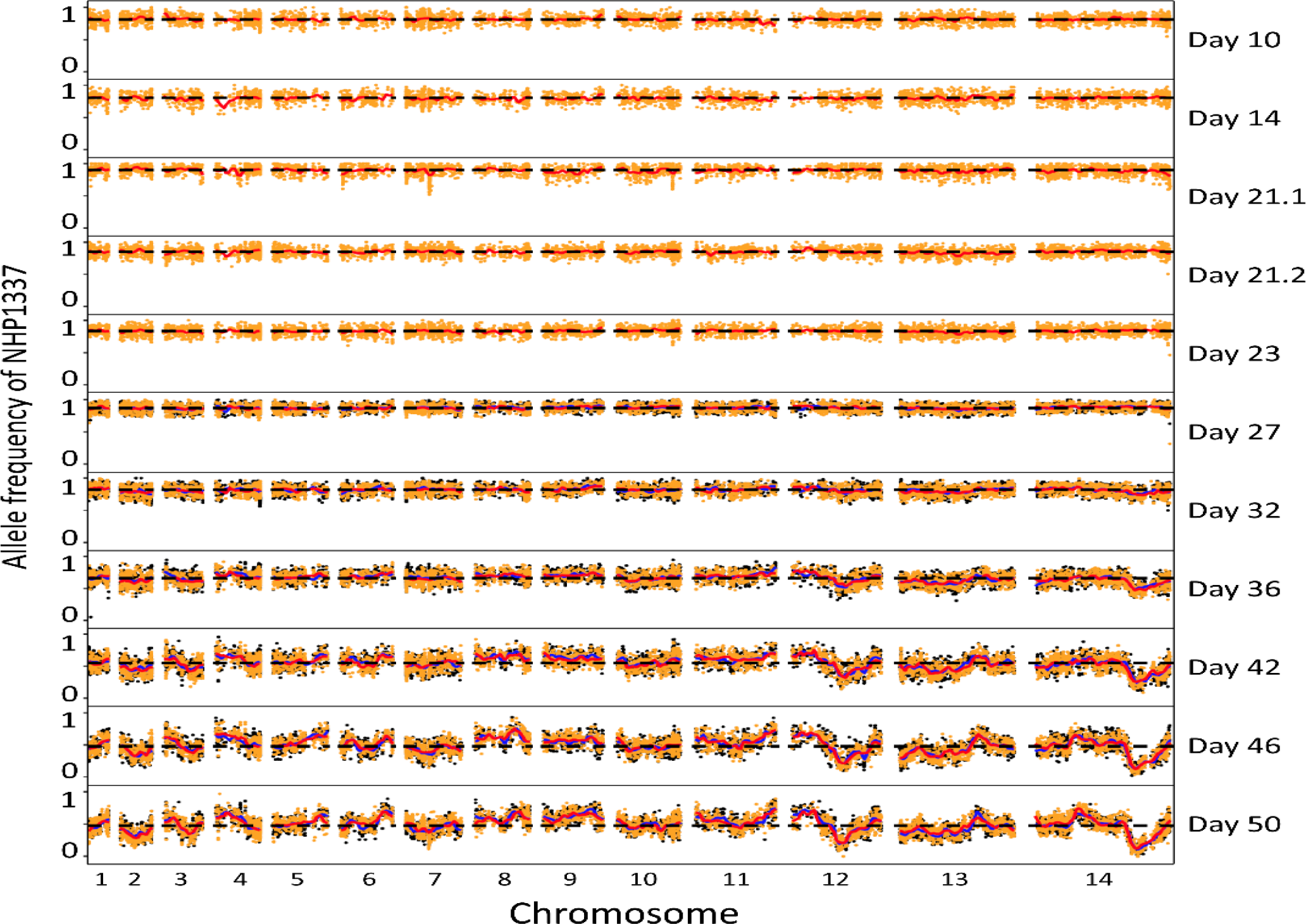
Plot of allele frequencies across the genome through *Plasmodium falciparum* life cycle. We divided the parasites into two replicates after two days’ *in vitro* culture (day 23). Orange and black indicated allele frequencies from these two parallel cultures. Red and blue lines showed tricube-smoothed allele frequencies. Black dashed lines indicated the average allele frequency across the genome. Sample collecting days were marked on the right. Day 10 shows allele frequencies of maturing oocysts, day 14 shows sporozoites, day 21.1 shows liver stage schizonts, day 21.2 shows transitioned blood stage parasites, and day 23-50 shows fifteen asexual cycles in blood stage culture.

**Figure 4.**
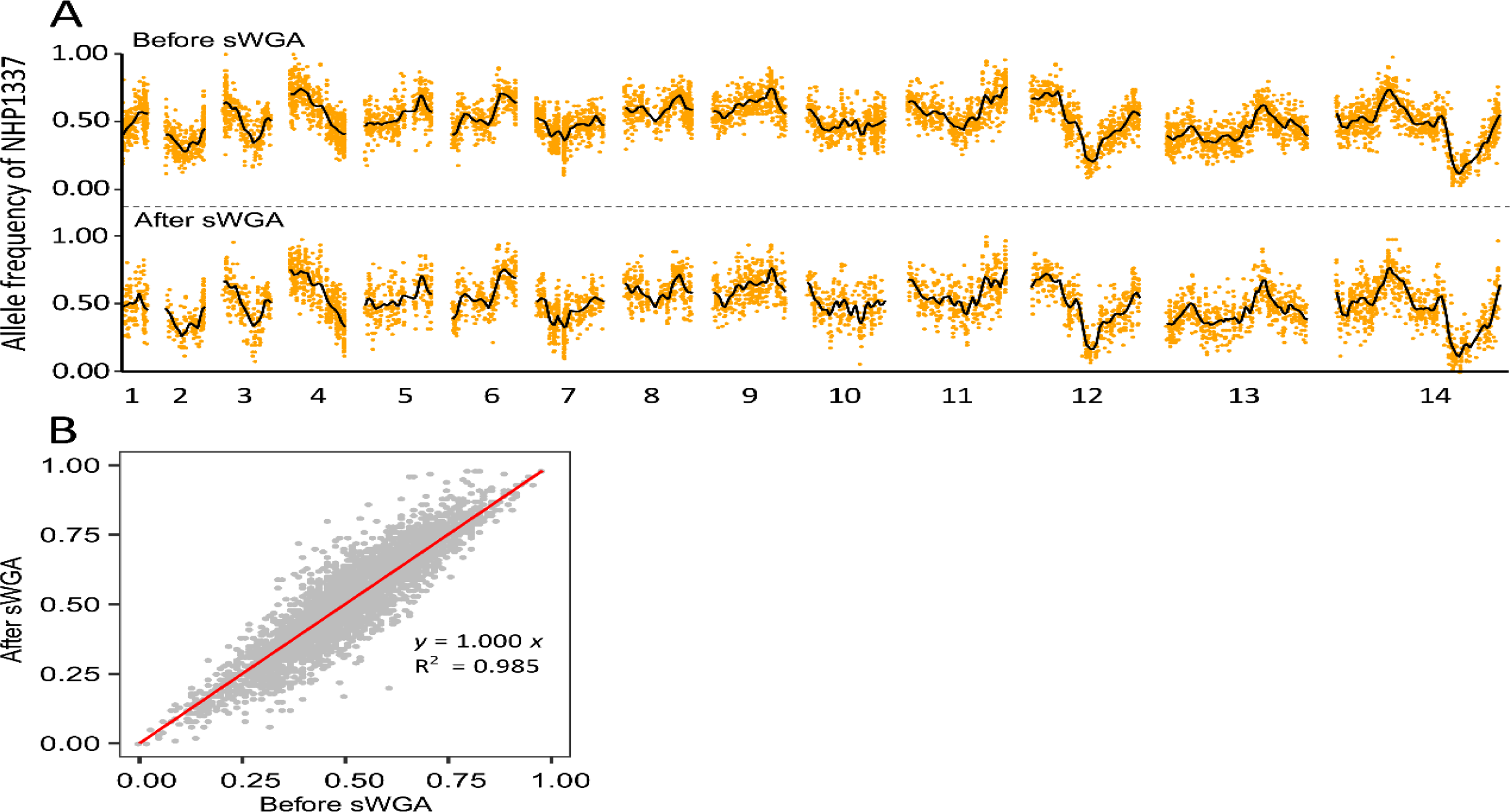
Allele frequencies estimated before and after selective whole genome amplification (sWGA). (A) Plot of allele frequencies across the genome. (B) Concordance between allele frequencies estimated before and after sWGA.

### Evaluation of bias in allele frequency measurement

To evaluate the accuracy of allele frequencies estimated after sWGA, we sequenced blood samples using both the sWGA-WGS approach and the WGS approach. We plotted allele frequencies of the parent NHP1337 across the genome and tricube-smoothed the frequency with window size of 100kb to smooth out noise and estimate changes in adjacent regions. With 10 million 150 bp pair-end sequencing reads, there were fewer loci detected with coverage > 30× by the sWGA-WGS approach relative to WGS (5,024 loci by sWGA-WGS and 7,844 loci by direct WGS). The allele frequency trends, however, were highly consistent after smoothing (Fig. 4A). The allele frequencies estimated before and after sWGA were strongly concordant (*R*^2^ = 0.985, Fig. 4B), which strongly supports the comparability of these two different methods.

### Allele frequency changes in segregant pools

a. Mosquito stages: *Plasmodium* sexual blood stage infections differentiate into both male and female gametes and mate; consequently, selfed progeny, resulting from the fusion of gametes from the same parasite genotype, can occur (i.e., NHP1337 male gametes fertilizing NHP1337 female gametes and MKK2835 male gametes fertilizing MKK2835 female gametes). Selection towards selfed progeny is evident from skewing and shifting of whole genome allele frequencies. To investigate population composition at different infection stages, we plotted the allele frequency distribution of *Plasmodium* mitochondria and across the core genome (Fig. 2). We observed a strong skew (>80%) towards alleles from the ART-R parent in the mosquito stages, which suggests that many selfed progeny from NHP1337 were present. We sequenced parasite progeny cloned from day 21 mouse blood, which indicated that this is the case (Button-Simons et al. in preparation).
b. Liver stage: The allele frequency in the progeny parasite population shifted significantly towards the ART-R parent (NHP1337) at the liver stage. This is evident from comparisons of allele frequency distributions in the liver with those from sporozoites (Fig. 2A, Cohen’s *d* test, large effect size = 0.89). This skew observed in the liver stage is reduced in merozoites emerging from the liver (Cohen’s *d* test, medium effect size = 0.61).
c. Blood stages: During *in vitro* culture, the allele frequency of NHP1337 (ART-R) dropped to 50%, between day 32 and day 40 (Cohen’s *d* test, effect size = 1.65). We maintained replicate *in vitro* blood cultures from day 23. Highly repeatable skews were observed in allele frequencies across the genome in these two parallel cultures (Fig. 2A&B, *R*^2^=0.985). Furthermore, we observed the same skews in both the mitochondria and across the core genome (Fig. 2C), strongly suggesting that the selection was against NHP1337 selfed progeny. The NHP1337 selfed progeny were almost eliminated by day 42 and we thus estimated the selection coefficients against the NHP1337 selfed progeny. We observed strong selection against NHP1337 alleles, with *s* = 0.24±0.02 in the core autosomal genome and *s* = 0.22±0.01 in mitochondria (Fig. 2D). There is no significant difference between these two estimates (*p* = 0.363).

### Loci under selection

To pinpoint the loci that determine parasite fitness at each life cycle stage, we first plotted the whole genome allele frequencies throughout the life cycle (Fig. 3). In addition to the whole genome skew described above, we also observed specific regions of the genome that showed distortion in allele frequency after day 32. The skews in allele frequencies were remarkably consistent between the two replicate blood stage cultures, suggesting pervasive selection at multiple loci across the genome. We calculated G’ values to measure the significance of allelic skews (Fig. 5A; Supplemental Fig. S3; Supplemental Table S1). Two strong QTLs were identified on chr 12 and 14, with genome-wide false discovery rate (FDR) < 0.01. We further used Δ (SNP-index) to determine the direction of the allele frequency changes (Fig. 5B; Supplemental Fig. S4). In both regions, alleles from NHP1337 (ART-R) were selected against. We then calculated selection coefficient (*s*) across the genome (Fig. 5C). We observed particularly strong selection at these two QTL regions, with *s* = 0.12 on chr12 and *s* = 0.18 on chr14. In addition, there were a set of lower confidence QTLs with lower allele frequency changes and less impact on parasite fitness uncovered across the genome (Fig. 5; Supplemental Table S1).

**Figure 5.**
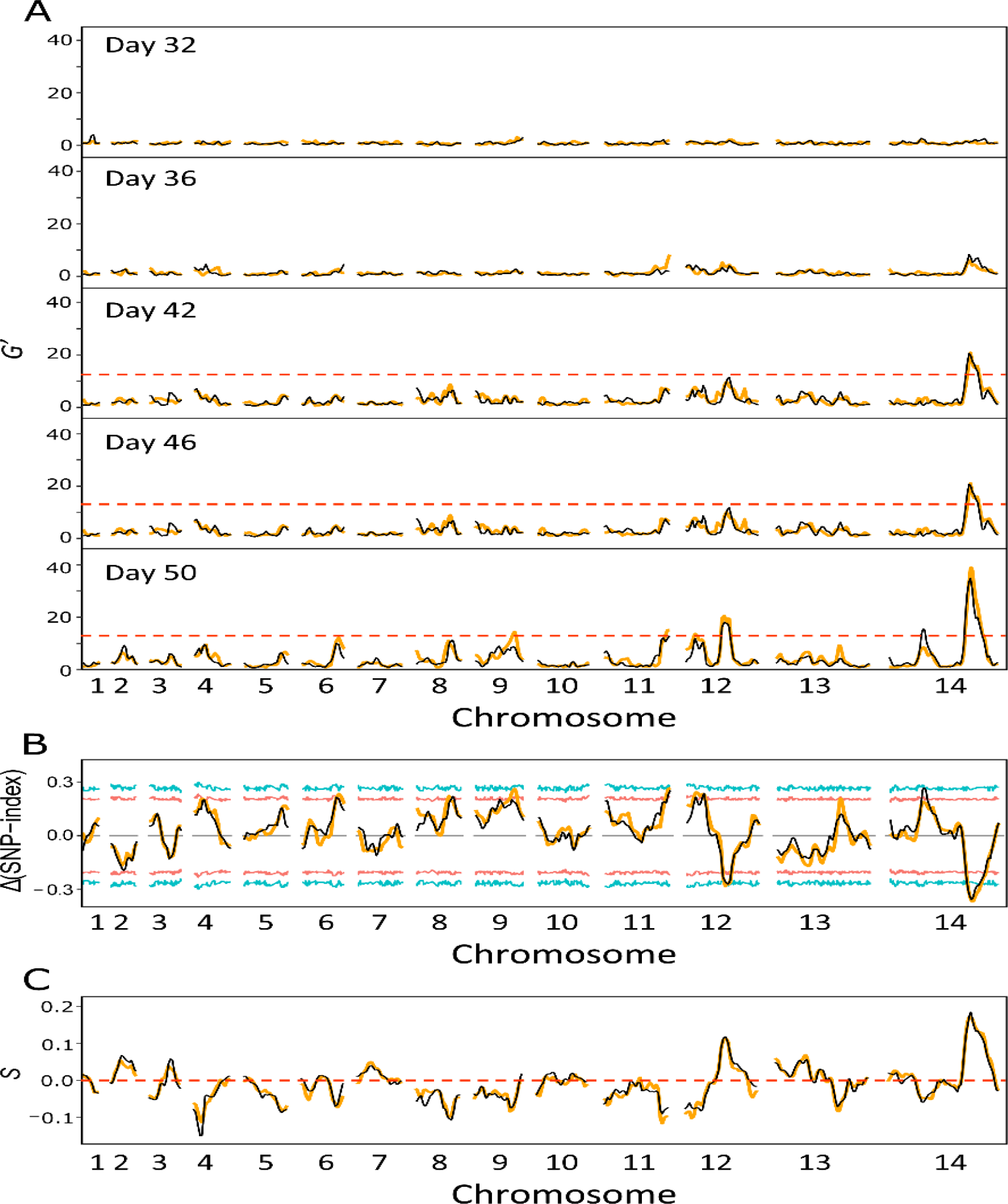
Bulk segregant analysis. (A) QTLs were defined with G’ approach by comparing allele frequencies at each loci to the average allele frequency across the genome. Regions with FDR > 0.01 were taking as significant QTLs. (B) Δ(SNP-index) for day50 progeny pools. The Δ(SNP-index) was the difference between SNP-index of each locus and the genome-wide average SNP-index. A positive Δ(SNP-index) value indicates an increase in alleles from NHP1337. Red and blue lines showed the 95% and 99% confidential intervals that matched with the relevant window depth at each SNP. (C) Tricube-smoothed selection coefficients (s). Estimation of *s* was based on the changes of allele frequency from day25 to day50. The mean selection coefficient was adjust to 0 to remove the influence of selfed progeny. Positive values of *s* indicate a disadvantage for alleles from NHP1337. Orange and black lines indicate experimental replicates.

### Fine mapping of chr 12 and 14 QTLs

We calculated 95% confidence intervals to narrow down the genes driving selection within the two QTL regions. The QTL on chr 12 ranged from 1,102,148 to 1,327,968 (226 kb) and the QTL on chr 14 ranged from 2,378,002 to 2,541,869 (164 kb).

#### Chr 12

The QTL region contained 48 genes, with 27 genes bearing at least one non-synonymous mutation differentiating the two parents (Fig. 6; Supplemental Table S2). Among the candidate genes with functional annotation, the multidrug resistance-associated protein 2 gene (*mrp2*, PF3D7_1229100) was located at the peak of the chr 12 QTL (Fig. 6A; Supplemental Table S2). The *mrp2* allele from NHP1337 carries three indels (3-24 bp) within coding microsatellite sequences compared with that in MKK2835. These indels don’t interrupt the open reading frame.

**Figure 6.**
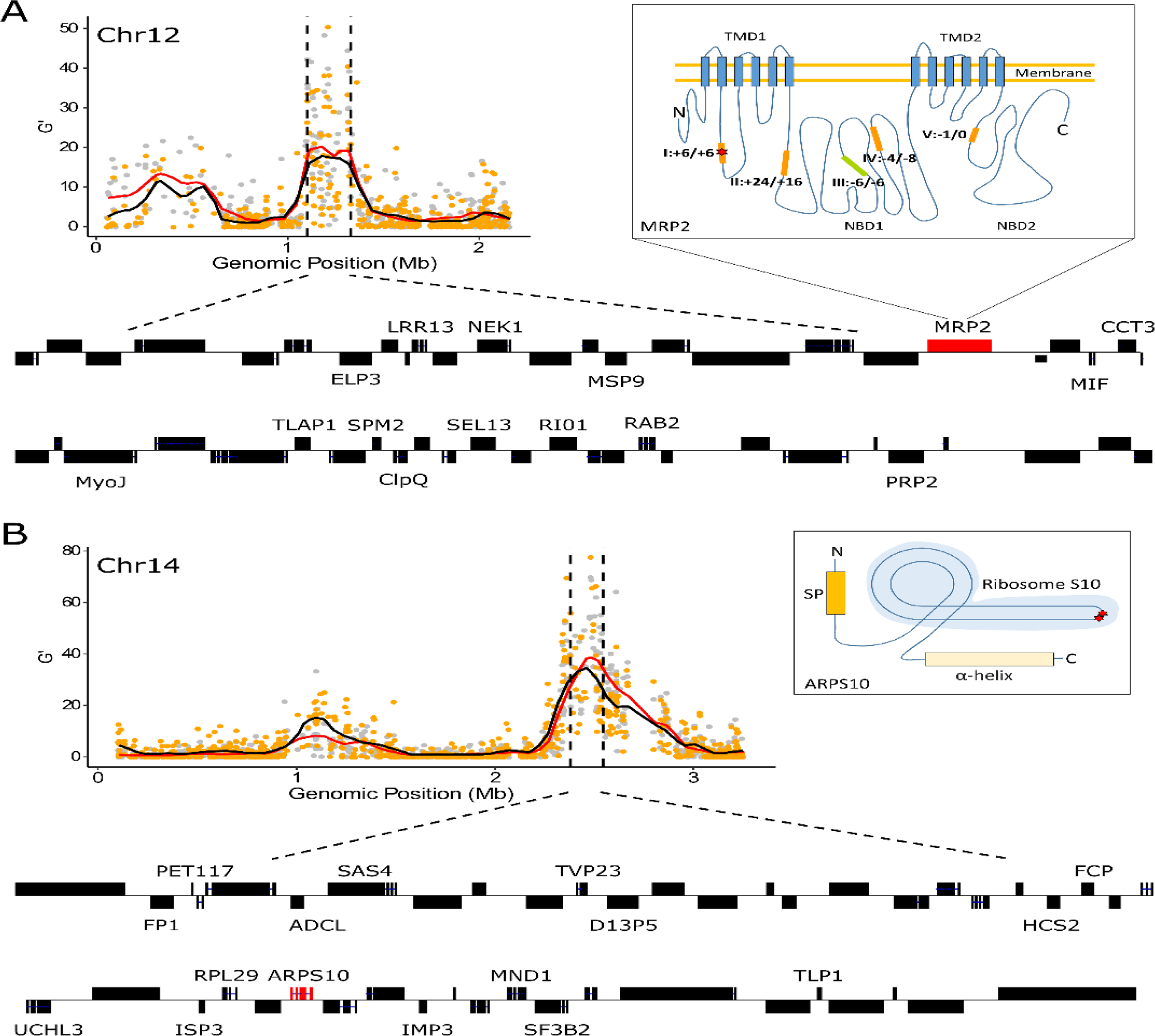
Overview of the genes inside of QTL regions on chr 12 (A) and chr 14 (B). Black dashed vertical lines are boundaries of the 95% confidential intervals (CIs) of the QTL. The QTL on chr 12 spanned 226 kb and included 48 genes, and the QTL on ch14 spanned 164 kb and included 45 genes. 2D structure of MRP2 and ARPS10 are presented in boxes next to the G’ plot. The structure of MRP2 was adapted from Velga et al., 2014. There are 5 microindels in the coding region of Pfmrp2 gene (I-V, orange and green blocks). Four of the microindels (orange blocks, 1 SNP and 3 indels) are different between ART-S and ART-R parental strains. The changes in peptide length relative to *P. falciparum* 3D7 are indicated next to the microindels, as microindel: ART-S/ART-R. ART-S and ART-R parasites have the same amino acid insertion at microindel I, but the sequence includes a synonymous mutation. The structure of ARPS10 is predicted by I-TASSER. ART-R has two non-synonymous mutations in gene *ARPS10*, Val127Met and Asp128His (red star). TMD: transmembrane domain; NBD: nucleotide-binding domain; SP: signal peptides.

### Chr 14

There are 45 genes located in this QTL and 13 contained non-synonymous mutations that distinguish the two parents (Fig. 6; Supplemental Table S2). The gene encoding apicoplast ribosomal protein S10 (*arps10*, PF3D7_1460900) was located at the peak of this QTL. There are two non-synonymous mutations (Val127Met and Asp128His) detected in *arps10* from NHP1337 as compared to MKK2835. The Val127Met mutation was suggested to provide a permissive genetic background for artemisinin resistance-associated mutations in *kelch13* in a genome-wide association analysis (Miotto et al. 2015).

## Discussion

### Pervasive selection in a *Plasmodium* genetic cross

In this experiment, we observed both genome-wide selection against selfed progeny, and locus specific selection that resulted in skews in the frequency of particular parental alleles in progeny.

#### (i) Genome-wide selection against selfed progeny

Initially, frequencies of alleles derived from the two parental parasites were strongly skewed (0.81 ± 0.08) towards the NHP1337 parent. This deviation from the expected 0.5 ratio for outcrossed progeny occurs because hermaphroditic malaria parasites produce both male and female gametocytes; fusion between male and female gametes of the same genotype (selfing) is possible. The simplest explanation for this observed skew is that an excess of selfed progeny were generated from the NHP1337 parent genotype compared to MKK2835 parent. We also dilution cloned progeny collected on day 23 of the experiment, which confirmed our suspicion that selfing of NHP1337 leads to the skew in allele frequency (Button-Simmons et al. in preparation). It is unclear whether the excess of selfed progeny from the NHP1337 parent relative to the MKK2835 parent results from an imbalance in gametocytes from these parental parasites when staging the cross, or from inherent differences in propensity to self in these two parasite clones.

Frequencies of NHP1337 remained high from day 10 (mature oocysts) until day 30 (after 10 days of *in vitro* blood culture). At this point, genome-wide frequencies of the NHP1337 parasite declined significantly from 0.85 to 0.54 on day 42. We observed a parallel decline of both mitochondrial and autosomal allele frequencies for the NHP1337 parasite. This is consistent with selection removing selfed NHP1337 genotypes from the progeny, otherwise we would expect selection on these two genomes to be decoupled. Selection was extremely strong (mitochondrial *s* = 0.22 ±0.01; autosomal *s* = 0.24 ±0.02) for both genomes. Furthermore, we observed the same patterns using whole genome sequencing and amplicon sequencing for measuring allele frequencies of mtDNA (Supplemental Fig. S2), suggesting that our results are robust to any methodological biases. Analysis of further crosses will allow us to determine whether selection against selfed progeny is a general feature of crosses in malaria.

These data demonstrate systematic selection against parental genotypes generated by selfing of the ART-R parent. Reproduction by outcrossing is prevalent in nature, even in hermaphroditic species (Charlesworth and Willis 2009). Inbreeding leads to reduced fitness of offspring (inbreeding depression), while outbreeding among genetically differentiated individuals improves the performance of the F1 generation (heterosis) (Whitlock et al. 2000; Charlesworth and Willis 2009). We observed strong selection against selfed NHP1337 genotypes which resulted in elimination of selfed progeny in six asexual cycles (day 30-42). Possible explanations for the lower fitness in selfed progeny are: (1) recombination allows removal of deleterious mutation in outcrossed progeny. Accumulation of deleterious mutations occurs during clonal expansion and in inbred parasite lineages. Both parental parasites used in this cross were isolated from Southeast Asia, an area of low parasite transmission intensity, where most infected patients harbor a single parasite genotype. As a consequence mosquito blood meals contain male and female sexual stages from the sample parasite clone, and therefore deleterious mutations can accumulate since inbreeding predominates (Anderson et al. 2000; Nkhoma et al. 2013). We speculate that recombinant genotypes generated by outcrossing between NHP1337 and MKK2835 parents have reduced numbers of deleterious alleles and therefore outcompete inbred parental genotypes. (2) *In vitro* culture, where the strongest selection was observed in this experiment, represents an ecological niche change for both parental genotypes. Recombinants generated by outbreeding may be more fit in these laboratory conditions.

There is an interesting shift in allele frequencies between sporozoites sampled from mosquito salivary glands and liver stage parasites recovered from infected mice on day 21 (Fig. 2), with liver stage parasites carrying high frequencies of NHP1337 alleles (liver 0.89 vs sporozoites 0.79, with large Cohen’s *d* effect size [0.89]). The allele frequency of parasites from *in vivo* blood collected on the same day is 0.84, which is between those from sporozoites and liver stage parasites. During liver stage development, single sporozoites take up residency within hepatocytes and divide mitotically over the course of ∼7 days (determined with laboratory strains of *P. falciparum* NF54 (Vaughan et al. 2012)) until liver schizonts burst releasing tens of thousands of merozoites into the blood. The simplest explanation of the observed allele frequency shift is a genotype-dependent variation in the duration of parasite liver stage development. We suggest that the selfed NHP1337 progeny remain in the liver longer and thus at the day 7 sampling, recombinant liver stage parasites have already transitioned to blood stage, generating the observed difference in allele frequencies. Further work is needed to directly determine the duration of liver stage development and if other liver stage parasite phenotypes (schizont size/merozoite numbers) differ among parasite genotypes.

#### (ii) Locus specific selection

We observe a progressive increase in the variance of allele frequencies of SNPs from day 30-50 (during blood stage culture) (Fig 2A). Several features of these data suggest that this is primarily driven by selection, rather than genetic drift. First, we note an extremely strong repeatability in allele frequency skews across the genome in the two replicate parasite cultures established from the humanized mouse infection. This is reflected in the high correlation between allele frequencies between these two replicates at the end of the experiment (Fig 3, day 50) when variance in allele frequency is at its maximum. The strong repeatability in patterns of skew observed suggest that there are multiple loci across the genome that influence parasite growth rate and competitive ability. Second, we see several regions of the genome that show extreme skew relative to the genome wide average. Two genome regions in particular (on chr 12 and on chr 14) show strong and significant skews that cannot be explained by drift. These allelic disproportions also increase progressively from 25-50 days, consistent with selection coefficients (*s*) of 0.18/48 hr asexual cycle for the chr 14 locus and (*s*) of 0.12/48 hr asexual generation for the chr 12 locus.

The strong selection observed against particular alleles segregating in this genetic cross (in the absence of drug pressure) is surprising, given that both parental parasites were isolated from patients. How can such strongly disadvantageous alleles be maintained in natural parasite populations? We suggest three explanations. First, we think that the most likely explanation is that the fitness of these alleles may depend on genetic background (Lynch 1991) and reflect epistatic interactions. We note that of the two parental strains used in this study, MKK2835 (ART-S) was isolated in 2003, while NHP1337 (ART-R), was collected in 2013. In the 10 years between 2003-13, artemisinin-resistant parasites spread to high frequency on the Thailand-Myanmar border (Anderson et al. 2016). Intense drug selection in this 10-year interval has led to accumulation of additional genetic changes associated with ART-R, which may act epistatically together with other ART-R-associated genes (Cerqueira et al. 2017). It is certainly interesting that the chr 14 QTL contains *arps10*, which has been suggested to provide a permissive background for ART-R evolution (Miotto et al. 2015). Outcrossing between individuals with different adaptations can result in disruption of this selective advantage, resulting in a loss of fitness (Coyne 2004). Second, there is a possibility that *de novo* deleterious mutations in these two QTL regions were fixed in the cloned NHP1337 parasites during the brief period of laboratory culture. We think this is unlikely because we also see pervasive selection at multiple genes outside these two major QTL regions, just with lower significance using G’ statistics. Third, we cannot discount the possibility that the strong selective disadvantages observed within these QTL regions reflects the artificial nature of this system with humanized mice and asexual culturing of parasites. During normal transmission in the field selection against these genes may not be present.

We anticipate that intensity of competition among parasite clones within infected patients may closely parallel the patterns we observe within our genetic cross. The estimated occurrence rate of mixed infections ranges from 18% to 63% in African and Southeast Asia countries (Anderson et al. 2000; Zhu et al. 2018). Although there was likely more intense competition in this experimental cross, with millions of sporozoites infecting a single mouse, single cell sequencing has revealed seventeen unique clones in a single human infection (Nkhoma et al. 2018), which suggests that similar competitive interactions will also occur in patients. We note that while the intensity of competition may be similar in humanized mice, *in vitro* parasite cultures or infected humans, the nature of selection may differ. In infected people, parasite genotypes that allow evasion of immunity or alter parasite cytoadhesion properties may be selected, while growth competition is likely to be the predominant selective force in immunosuppressed humanized mice or *in vitro* culture.

#### (iii) What drives QTL peaks on chr12 & chr14?

Inspection of the genes under the QTL peaks allows us to speculate about specific genes that may be driving the selection observed by Miotto et al.(2015) showed that four different non-*kelch13* loci (ferredoxin, *fd*; apicoplast ribosomal protein S10, *arps10*; multidrug resistance protein 2, *mdr2*; chloroquine resistance transporter, *crt*) are associated with the resistance phenotype, but not directly responsible for resistance. They suggested that a suite of background mutations was a prerequisite for mutations in *kelch13*. In our experiment, *arps10* falls near the peak of the strongly selected chr 14 loci (Fig 6), which could suggest a functional relationship. We examined the presence of the background mutations found in both parental strains. The ART-R parent, NHP1337, contains mutations in all of four of the genes described by Miotto et al. (Miotto et al. 2015) (*fd*, *mdr2, crt and arps10*), while the ART-S parent, MKK2835, contains just three of these mutations (*fd*, *mdr2* and *crt*), so only *arps10* mutations are segregating in this cross. It will be interesting to test the role of the remaining three loci (*fd*, *mdr2* and *crt*) by conducting additional experimental crosses.

The multidrug resistance-associated proteins (MRPs), belong to the C-family of ATP binding cassette (ABC) transport proteins that are well known for their role in multidrug resistance. Rodent malaria parasites encode one single MRP protein, whereas *P. falciparum* encodes two: MRP1 and MRP2 (Rijpma et al. 2016). Several studies have shown that PfMRP1 is associated with *P. falciparum’s* response to multiple anti-malaria drugs and that disruption of PfMRP1 influences the fitness of parasites under normal culture conditions (Mu et al. 2003; Dahlström et al. 2009; Raj et al. 2009). The function of PfMRP2 is not as well understood. Transfection studies have shown that MRP2-deficient malaria parasites are not able to maintain a successful liver stage infection (Rijpma et al. 2016; van der Velden et al. 2016). In our study, *mrp2* was found located at the peak QTL on chr12. We speculate that *mrp2* may also play a role in parasite fitness during asexual parasite stages. However, we cannot exclude that other neighboring loci may drive the observed allele frequency changes.

### No selection against the *kelch13*-C580Y allele conferring ART resistance

Interestingly, we do not see evidence for selection against the *kelch13*-C580Y allele (chr 13) that underlies resistance to ART treatment. We previously used CRISPR/Cas9 editing to add the C580Y substitution to a wild type parasite (Nair et al. 2018). Head-to-head competition experiments revealed strong fitness costs (*s* = 0.15/asexual cycle) associated with this substitution. In agreement, Straimer et al (Straimer et al. 2017) conducted similar experiments with Cambodian parasites: they showed that the addition of the C580Y resulted in strong fitness costs for some parasites, but had no fitness impact in recently isolated Cambodian parasites. These data also suggest that epistatic interactions with other loci may compensate and restore parasite fitness. We suspect that this may also be the case in our experiment.

### Technical considerations & caveats

#### Maximizing statistical power

Our statistical power to detect QTLs is limited by the number of recombinants generated. In our experiment, the mouse was infected with 204 mosquitoes carrying on average three oocysts. Given that each oocyst is expected to contain sporozoites representing up to four different genotypes (i.e. a tetrad), the maximum number of unique recombinants is 204 × 3 × 4 = 2448 in this cross. We can increase the power of these experiments using mosquitoes with higher infection rates. We routinely obtain an average of 10 oocysts/mosquito, so can potentially increase numbers of recombinants by at least three-fold with the same number of mosquitoes. A second advantage of humanized mice over splenectomized chimpanzees as an infection model is that we can easily increase numbers of humanized mice used per cross. By using independent pools of mosquitoes to infect mice, we can multiply the numbers of recombinants generated, while also establishing true biological replicates of each experiment.

In this experiment, we found large numbers of inbred progeny generated by mating between male and female gametes of the same genotype. While we were able to use these to document selection against selfed progeny, this reduces the number of recombinant progeny and therefore limits statistical power for locating QTLs. A method that maximizes outcrossing would be particularly useful for future crosses. For example, aphidicolin treatment has been successfully used in rodent malaria systems to kill male gametes (Ramiro et al. 2015).

#### Combining BSA with cloning recombinant progeny to detect epistasis

BSA cannot be used to directly examine epistatic interactions, due to the lack of haplotype information. Fortunately, *P. falciparum* has a key advantage over rodent malaria systems because parasites can be grown *in vitro* and cloned by limiting dilution. Hence, BSA can be complemented by cloning progeny from the same genetic cross and directly examining haplotypes carrying different allele combinations. Furthermore, we can use BSA to directly test for interactions between genes. For example, we suspect that interactions between *kelch13* mutations and *arps10* may drive the skew observed at chr 14. This hypothesis can be directly tested by repeating the cross with parasites that have been edited to remove the *kelch13* mutation or candidate *arps10* mutations, to see if the skew on chr 14 disappears.

#### sWGA performance

The sWGA method efficiently enriches *P. falciparum* DNA from infected mosquito and mouse tissues, confirming the performance of this approach for enriching parasite DNA from dried blood spots (Guggisberg et al. 2016; Oyola et al. 2016; Sundararaman et al. 2016; Cowell et al. 2017). Our results further show that sWGA does not generate bias in allele frequency measurement (Fig. 4). However, sWGA does have limitations with highly contaminated samples, such as early infected mosquitoes (four days post infection). DNA extracted from day 4 midguts typically contains > 99.99% mosquito DNA. Only 4.3% of sWGA products from these samples was *Plasmodium* DNA. In contrast, we were able to obtain > 88% of parasite DNA from sWGA, with starting material containing ≥ 1% *P. falciparum* DNA (Table 1).

#### Potential of BSA for examining selection in the mosquito stage

We did not observe allele frequency changes during mosquito infections in this experiment. We suggest two reasons for this. First, the *A. stephensi* mosquito used is originally from urban India and widely spread across Southeast Asia, and therefore may show good compatibility with Southeast Asian parasites. Furthermore, this specific mosquito line has been long-term lab adapted, and is highly susceptible to infection with multiple parasite lines. Second, the infection period in mosquitoes in this experiment is relatively short, because we sacrificed all the mosquitos in two weeks. As a consequence, we can only detect very strong selection at this stage. However, hard selection resulting from incompatibility between parasites and mosquitoes should still be possible to detect and map in this system. We note that Molina-Cruz et al (Molina-Cruz et al. 2013) were able to determine parasite QTLs for compatibility between *P. falciparum* and mosquitoes using parasite progeny derived from the original malaria crosses conducted in chimpanzees, providing proof-of-principal that this is possible.

### Future prospects

Human malaria can now be passaged through humanized mice, as well as grown *in vitro* in culture and cloned. The power of the BSA approach has been clearly demonstrated in rodent malaria, where it has been used to identify the genetic components controlling a broad range of selectable phenotypes, including virulence and immunity, growth rate and drug resistance (Rosario et al. 1978; Culleton et al. 2005; Martinelli et al. 2005; Pattaradilokrat et al. 2009; Hunt et al. 2010). However, human malaria parasites and rodent malaria parasites are genetically distant and human parasites show numerous unique biological features not found in rodent malaria parasites. Our approach can now be applied to directly study multiple selectable traits in the human parasite *P. falciparum* via genetic crosses. We anticipate that BSA will provide a powerful complement for the study of *P. falciparum* genetics.

## Material and Methods

### Ethics statement

The study was performed in strict accordance with the recommendations in the Guide for the Care and Use of Laboratory Animals of the National Institutes of Health (NIH), USA. To this end, the Seattle Children’s Research Institute (SCRI) has an Assurance from the Public Health Service (PHS) through the Office of Laboratory Animal Welfare (OLAW) for work approved by its Institutional Animal Care and Use Committee (IACUC). All of the work carried out in this study was specifically reviewed and approved by the SCRI IACUC.

### Parasites, mosquitoes and mice

FRG NOD huHep mice (Azuma et al. 2007) with human chimeric livers were purchased from Yecuris Corporation. Mice used in the study were supplemented with NTBC at 8 mg/L in their drinking water on arrival and maintained on this dose until euthanasia. The *A. stephensi* mosquitoes used in this study were maintained at 27 °C and 75% humidity on a 12-h light/dark cycle. We followed MR4 protocols (Moll et al. 2008) for larval stage and adult stage rearing; larvae were fed with finely ground Tetramin fish food, and adults were fed with cotton balls soaked in a solution of 8% dextrose and 0.05% para-aminobenzoic acid in water.

*P. falciparum* blood stage cultures were maintained *in vitro* in standard cell culture media (RPMI-1640 with 25 mM HEPES and 2 mM l-glutamine supplemented with 50 μM hypoxanthine and 10% A+ human serum) (Moll et al. 2008). An atmosphere of 5% CO_2_, 5% O_2_ and 90% N_2_ was used for growth, and infected *P. falciparum* red blood cells were subcultured into O+ erythrocytes.

We used two parasites isolated from hyper-parasitemic patients visited the Wang Pha clinic run by the Shoklo Malaria Research Unit (SMRU). *P. falciparum* NHP1337 (ART-R, C580Y mutant *kelch13*) and MKK2835 (ART-S, wild-type *kelch13*) isolates were grown in the laboratory, cloned by limiting dilution, and single parasite clones were used for these experiments.

### Preparation of genetic cross and sample collection

We generated the cross as described by Vaughan *et al.* (Vaughan et al. 2015). We collected samples from infected mosquito midgut and salivary gland, mouse liver and *in vivo* blood, and *in vitro* blood cultures (Fig. 1). To initial the cross, we set up asexual cultures of both parents by at 1% (mixed stages) parasitemia and 5% hematocrit. The cultures were maintained with daily medium changes for two weeks to enrich gametocytes. We then mixed gametocytes from each parent at equal ratio, and fed to adult female mosquitoes. This day was defined as day 0 for sample collecting (Fig. 1; Table 1). Forty-eight midguts were dissected at each oocyst collection time point (day 4 and day 10). The prevalence of infection was analyzed at day 10. Salivary gland were separated to collect sporozoites at day 14 after infection. Sporozoites from 204 mosquitoes were mixed together for infection into the mouse and for isolation of genomic DNA.

Six days after sporozoite injection (day 20), we injected mice intravenously with 400 μL of packed O+ huRBCs. The intravenous injection was repeated the next day (day 21). Four hours after the second huRBC injection, mice were sacrificed and blood was removed by cardiac puncture in order to recover *P. falciparum*–infected huRBCs. The mouse liver was dissected, immediately frozen in liquid nitrogen and then stored at −80 °C. The blood was added to 10 mL complete medium (RPMI-1640 with 25 mM HEPES, 2 mM l-glutamine, and 50 μM hypoxanthine) and pelleted by centrifugation at 200g. We then removed the supernatant along with the buffy coat (containing white blood cells), and the red blood cells were washed three times with 10 mL complete medium, with pelleting and centrifugation as detailed above. After the third wash, an equal volume of packed O+ huRBCs (approximately 400 μL) was added, and the total RBC pellet was resuspended in complete medium to 2% hematocrit, and maintained in an atmosphere of 5% CO_2_, 5% O_2_ and 90% N_2_. Two days after culture, the parasites were split equally into two wells (repeat A and repeat B) of a standard six-well plate, and 50 μL of freshly packed huRBCs were added every 2 days to each well. Once parasitemia reached 4%, serial dilutions of parasites were carried out to maintain healthy cultures. The cultures were maintained for 30 days in total (day21-day50), and 50ul packed red blood cells (RBCs) were collected and frozen down every 2-4 days.

### DNA isolation and parasite DNA quantification

We extracted and purified genomic DNA using the Qiagen DNA mini kit, and quantified amounts using Qubit. We performed real-time quantitative PCR (qPCR) reactions to estimate the proportion of parasite genomes in each DNA sample. Reactions were performed in duplicate using the AB1 prism 7900HT Sequence Detection system (Applied Biosystems, Carlsbad, California, USA) as follows: 95 °C for 10 min, then 40 cycles of 95 °C for 15 s and 60 °C for 1 min. Duplicate reactions showing a difference in CT greater than one were rerun. We examined the melting curve (60–95 °C) at the end of each assay to verify the uniqueness of the PCR products generated. The reaction mixture consisted of 5 µL of SYBR Green MasterMix (Applied Biosystems, Carlsbad, California, USA), 0.3 µL of 10 µM primers (Table S3) amplifying 67 bp of the *Pf_Tubilin* gene (PF3D7_1008700), 3.4 µL of sterile water and 1 µL of total DNA template. We plotted standard curves using seven dilutions at copies/µL from 2 × 10^1^ to 2 × 10^7^ with 10-fold interval of a purified *Pf_Tubilin* PCR product. The number of *Pf_Tubilin* copies in each sample was estimated according to the standard curve.

### Selective whole genome amplification

We used selective whole genome amplification (sWGA) to enrich parasite DNA for samples obtained from infected mosquito and mouse tissues. sWGA reactions were performed following Oyola et al (Oyola et al. 2016). Each reaction (50 μl total volume) contained at least 0.2 × 10^6^ copies of *Plasmodium* DNA, 1× BSA (New England Biolabs), 1 mM dNTPs (New England Biolabs), 3.5 μM of each amplification primer (Oyola et al. 2016), 1× Phi29 reaction buffer (New England Biolabs), and 30 units of Phi29 polymerase (New England Biolabs). We used a PCR machine (SimpliAmp, Applied Biosystems) programmed to run a “stepdown” protocol: 35 °C for 10 min, 34 °C for 10 min, 33 °C for 10 min, 32 °C for 10 min, 31 °C for 10 min, 30 °C for 3 h then heating at 65 °C for 10 min to inactivate the enzymes prior to cooling to 4 °C. Sample were cleaned with AMPure XP Beads (Beckman Coulter), at a 1:1 ratio. We quantified the amplified product using Qubit^®^ dsDNA Broad Range (Thermo Fisher Scientific) to determine whether there was enough material for sequencing-minimum required is 50 ng. The sWGA products were further quantified by qPCR (described above) to confirm that the majority of the products were from *Plasmodium*.

### Whole genotype sequencing and mapping

We constructed next generation sequencing libraries using 50ng DNA or sWGA product following the KAPA HyperPlus Kit protocol with 3-cycle of PCR. All libraries were sequenced to an average coverage of 100x using an Illumina NEXTseq 500 sequencer.

We individually mapped whole-genome sequencing reads for each library against the *P. falciparum* 3D7 reference genome (PlasmoDB, release32) using the alignment algorithm BWA mem (http://bio-bwa.sourceforge.net/) under the default parameters. The resulting alignments were then converted to SAM format, sorted to BAM format, and deduplicated using picard tools v2.0.1 (http://broadinstitute.github.io/picard/).

### Mitochondrial DNA Amplicon sequencing

We used amplicon sequencing to trace the biases in mtDNA transmission, as sWGA with circular DNA may swamp out other sWGA products. We use at least 1000 copies of parasite genome as template for each reaction. Illumina adapters and index sequences were added to the PCR primers (Table S3). We cleaned PCR products with AMPure XP beads, and quantified products using a standard picogreen assay that can be read on a fluorescent plate reader. Equal number of molecules were pooled from each reaction and sequenced on Illumina NEXTseq 500 sequencer.

### SNP calling between parents

We first genotyped the two parental strains. We used Genome Analysis Toolkit GATK v3.7 (https://software.broadinstitute.org/gatk/) to recalibrate the base quality score based on a set of verified known variants (Miles et al. 2016). We called variants for each parent using HaplotypeCaller and then merged using GenotypeGVCFs with default parameters except for sample ploidy 1. We applied filters to the original GATK genotypes using quality criteria QD > 2, FS < 60, MQ > 40, SOR < 3, GQ > 50 and DP ≥ 3 for SNPs, and QD > 2.0, FS < 200, SOR < 10, GQ > 50 and DP ≥ 3 for indels. The recalibrated variant quality scores (VQSR) were calculated by comparing the raw variant distribution with the known and verified *Plasmodium* variant dataset. Loci with VQSR less than 1 were removed from further analysis.

### Bulk segregant analysis

We generated a “mock” genome using GATK *FastaAlternateReferenceMaker* from the genotype of parent NHP1337 (C580Y). The reads from bulk populations obtained at each stage of the lifecycle were mapped to this genome. Only loci with coverage > 30x were used for bulk segregant analysis (BSA). We counted reads with genotypes of each parent and calculated allele frequencies at each variable locus. Allele frequencies of NHP1337 were plotted across the genome, and outliers were removed following Hampel’s rule (Davies and Gather 1993) with a window size of 100 loci (Fig. 3).

We performed the BSA analyses using the R package QTLseqr (Mansfeld and Grumet 2018). We first defined extreme-QTLs by looking for regions with false discovery rate (FDR) < 0.01 using the G’ approach (Magwene et al. 2011). We then calculated the Δ (SNP-index) to show the direction of the selection (Takagi et al. 2013). Once a QTL was detected, we calculated and approximate 95% confidence interval using Li’s method (Li 2011) to localize causative genes.

We also measured the fitness cost at each mutation by fitting a linear model between the natural log of the allele ratio (freq[allele1]/freq[allele2]) against time (measured in 48hr parasite asexual cycles). The slope provides a measure of the selection coefficient (*s*) driving each mutation (Dykhuizen and Hartl 1980). The raw *s* values were tricube-smoothed with a window size of 100 kb to remove noise (Nadaraya 1964; Watson 1964). A positive value of *s* indicates selection against alleles from the ART-R parent (NHP1337), while a negative value of s indicates selection for NHP1337 alleles.

### Data access

Raw sequencing data have been submitted to the NABI Sequence Read Archive (SRA, https://www.ncbi.nlm.nih.gov/sra) under the project number of PRJNA524855.

## Acknowledgment

This work was supported by an NIH program project grant P01 AI127338 (MF) and by NIH grant R37 AI048071 (TJCA). Work at Texas Biomedical Research Institute was conducted in facilities constructed with support from Research Facilities Improvement Program grant C06 RR013556 from the National Center for Research Resources. SMRU is part of the Mahidol Oxford University Research Unit supported by the Wellcome Trust of Great Britain.

